# Chagas disease awareness and socioeconomic characteristics of Bolivian immigrants living in São Paulo, Brazil

**DOI:** 10.1101/775742

**Authors:** Rubens Antonio da Silva, Dalva Marli Valério Wanderley, Colin Forsyth, Ruth Moreira Leite, Expedito José de Albuquerque Luna, Nivaldo Carneiro Júnior, Maria Aparecida Shikanai-Yasuda, Grupo de Pesquisa em Doença de Chagas: atenção primária e imigração (Chagas Disease Research Group: Primary Care and Immigration)

## Abstract

In this study, part of a research project on Chagas disease among residents of Bolivia in São Paulo, we describe socioeconomic characteristics, knowledge about the disease and access to health services. A structured questionnaire was applied to a sample of 472 Bolivian adults (> 18 years) living in São Paulo enrolled in the Barra Funda School Health Center. The median age of participants was 28.5 years, 75.0% from the Bolivian department of La Paz, who were living in São Paulo for an average of 5.8 years. Regarding knowledge about the disease and exposure to certain risk factors, 47.7% indicated familiarity with the vector, 23.9% had seen vinchuca in their homes in Bolivia and 6.4% reported having been bitten by a triatomine. The conditions of living in rural areas in Bolivia or in other department than La Paz, have a relative with illness, high school graduation and have seen or been bitten by a vinchuca were significantly associated with the knowledge of the vector. This study provides a view on migration that has important implications for the distribution of Chagas’ disease and access to health care by providing subsidies for proposing public health policies.

**Author summary:** This article expresses part of the results of a research project called “Chagas disease in a population of Bolivian immigrants in São Paulo: an analysis of the prevalence of *Trypanosoma cruzi* infection and morbidity of Chagas disease, knowledge of the population about the disease and access to different levels of health care”. The problem of chronic Chagas disease occurs in many countries, including those not considered endemic, as a result of population movements, mainly by immigration due to urbanization which has led to its globalization. It is now considered an emerging disease with significant potential for transmission via blood transfusions, organ transplants and congenital via, in the absence of appropriate strategies in terms of public health, as well as reactivation of chronic disease in urban centers. It’s no different this phenomenon to the city of Sao Paulo. This study analyzed the sociodemographic inserts, labor, migration and knowledge about Chagas disease and its impact on personal, family and professional life of Bolivian immigrants living in São Paulo.

## Introduction

Chagas disease (CD) causes a greater burden of morbimortality than any other parasitic infection in the Americas, yet remains one of the world’s most neglected diseases[1]. Although regional collaborations such as the Southern Cone Initiative have made tremendous progress in curbing vector transmission [2], this success has not been replicated in the healthcare sphere; <1% of the over 6 million people living with the disease in the Americas [3] have been diagnosed and treated. The intense movement of human populations has been a defining feature of the past three decades and has profoundly impacted the epidemiology of Chagas and other diseases. While CD was traditionally confined to rural, endemic areas in Latin America, large numbers of affected people, spurred by political, economic, and environmental factors, have moved into nonendemic areas, including major urban centers of Latin America, and across national borders [4–5].

Much research has focused on the globalization of CD. In Europe, an estimated 68,000-122,000 people of Latin American origin live with CD, and another 326-347,000 live in the U.S. [6–7]. The demographic profiles are distinct, with the majority of European cases originating from Bolivia, while in the U.S. the highest numbers are from Mexico and Central America. A phenomenon which has received less attention is regional transnational migration within Latin America, its impact on CD epidemiology, and the implications for public health strategies. For instance, some research suggests Bolivian migrants in Buenos Aires may represent a group with higher risk for CD [8].

Migrants with CD confront unique challenges to accessing healthcare; they may possess limited economic resources; be excluded from local healthcare services; and experience difficulties communicating with providers due to linguistic, class and cultural barriers. Prior research has found a high degree of socioeconomic marginalization among immigrants with CD in Europe and the United States [9–10]. Despite having a higher risk for CD, immigrants from Latin America are typically undiagnosed and may not have heard of the disease [11–12]. Compounding this, providers in host countries are often unfamiliar with CD and not up-to-date with treatment recommendations [13–14]. These many barriers can prevent immigrants from accessing timely diagnosis and etiological treatment.

Brazil exemplifies many of the changing dynamics in CD’s social and epidemiological profile. In 2006, the Pan American Health Organization certified interruption of transmission by the main vector, *Triatoma infestans* [15]. Still, the WHO estimates over one million people infected with T. cruzi, while other estimates are much higher [2, 16]. Only a handful have received etiological treatment. Oral transmission is an emerging phenomenon [17], and internal migration has made CD a public health challenge in Brazil’s cities. São Paulo, the fifth largest city in the world, is also a major destination for transnational migration. Over 350,000 Bolivian immigrants live in the city [18] and a recent seroprevalence study in this population found 4.4% with *T. cruzi* infection [19]. Bolivians primarily migrate to São Paulo in search of job opportunities and greater economic stability for their families [20–22]. Prior research on Bolivian immigrants living near downtown São Paulo indicated a predominantly young population working mainly in garment sweatshops; the majority earned between 1-3 times minimum wage [23].

In this study, which forms part of a larger research project on CD among Bolivian residents of São Paulo, we describe socioeconomic characteristics, awareness of CD, and access to health services for women of chilbearing age and children in a sample of Bolivian immigrants. The goal is to gain knowledge and insight for strengthening healthcare policies and service delivery to help overcome the various barriers to diagnosis and treatment for CD which confront this migrant population.

## Methods

We administered a structured questionnaire to a sample of 472 Bolivian adults (>18 years old) living in São Paulo who were registered with the Dr. Alexandre Vranjac Escola Barra Funda Health Center (CSEBF is its Portuguese acronym; Escola Barra Funda refers to the neighborhood), a primary care clinic which is part of the Brazilian Public Health System. The CSEBF provides free primary healthcare services to individuals regardless of immigration status. From July to November 2013, participants were recruited while attending appointments at the CSEBF for various reasons. Participants who agreed to participate in the study underwent an informed consent process in Spanish. Roughly 95% of the patients who were approached agreed to participate.

The investigators created and utilized a Spanish-language questionnaire with multiple choice questions on sex, age, time living in São Paulo, education, income, employment situation, and other socioeconomic variables. Participants were also asked about familiarity with the “vinchuca” (the common name for the triatomine vector in Bolivia) and with CD, including its transmission and symptoms. Other questions focused on risk factors for CD, and for women of childbearing age, access to Brazilian maternal/reproductive health services. After asking for patients’ consent, Spanish-speaking interviewers administered the questionnaire. All participants were tested for CD.

Questionnaire data was coded and stored in a database for subsequent analysis. Statistical analyses were performed using Epidat v. 3.5.1 (Dirección Xeral de Saúde Pública, Galicia, Spain) and SPSS v. 25.0. (IBM, Armonk, NY, USA). In an initial univariate analysis, we calculated proportions and used chi-square tests to identify factors associated with awareness of CD; p≤ 0.05 was considered significant. We then used a multivariate logistic regression model to identify variables independently associated with knowledge of the vinchuca.

The study forms part of a larger research project, “Chagas disease in the Bolivian population of São Paulo: An analysis of prevalence of *Trypanosoma cruzi* infection, morbidity, knowledge, and access to healthcare”, which was financed by the National Council for Scientific and Technological Development-Department of Science and Technology CNPq / DECIT, Ministry of Health, Brazil (grant # 404336/2012). Ethical approval was obtained by the Committee of Ethics in Research at the Clinical Hospital of the School of Medicine of the University of São Paulo, written informed consent was requested from participants.

## Results

Participants’ median age was 28.5 years (Table 1), and three quarters came from the Bolivian department of La Paz. Respondents had lived in São Paulo for a median of 5.8 years. Most respondents (55.4%) had lived in São Paulo between 1-5 years, though 44% indicated >5 years’ residence. There were slightly more females than males, and more than half of respondents fell into the 18-29 age range, while <15% were over 40. Two thirds of respondents had completed a high school education. Nearly all respondents (99.0%) lived in two neighborhoods, Bom Retiro and Barra Funda. Over 90% worked in the garment industry. While 21.6% of participants reported per-person family income less than 678 R$ ($299US), the minimum wage at the time [24] over 50% earned between 1-2 times the minimum wage. Neither time in São Paulo nor education level were associated with significant differences in income.

**Table 1.** Sociodemographic characteristics of participants, study of Chagas disease awareness among Bolivian migrants in São Paulo

| Category | N |  | % |
| --- | --- | --- | --- |
| **Sex** |  |  |  |
| M | 217 |  | 45.9 |
| F | 255 |  | 54.1 |
| **Age group** |  |  |  |
| 18 – 29 | 251 |  | 53.2 |
| 30 – 39 | 154 |  | 32.6 |
| > 40 | 67 |  | 14.2 |
| **Years of residence in São Paulo** |  |  |  |
| < 1 | 11 |  | 2.3 |
| 1 – 5 | 252 |  | 53.4 |
| > 5 | 209 |  | 44.3 |
| **Lives with:** |  |  |  |
| Family | 216 |  | 45.8 |
| Relatives | 135 |  | 28.6 |
| Friends | 106 |  | 22.4 |
| Other | 15 |  | 3.2 |
| **Marital status** |  |  |  |
| Single | 143 |  | 30.2 |
| Married | 155 |  | 32.8 |
| Stable relationship | 161 |  | 34.1 |
| Other | 13 |  | 2.9 |
| **Occupation** |  |  |  |
| Garment industry | 428 |  | 90.6 |
| Other | 44 |  | 9.4 |
| **Family income** |  |  |  |
| ≤ one minimum salary | 102 |  | 21,6 |
| 1-2 times minimum salary | 250 |  | 52,9 |
| >2 times minimum salary | 120 |  | 25,5 |
| **Education level** |  |  |  |
| Primary or less | 15 |  | 3.2 |
| Some secondary | 137 |  | 29.0 |
| High school graduate | 298 |  | 63.1 |
| College | 22 |  | 4.7 |

We assessed awareness of CD and exposure to certain risk factors (Table 2). Nearly half (225/472, 47.7%) of participants indicated familiarity with the vector (*vinchuca*); 113 (23.9%) had seen the vinchuca in their homes in Bolivia, and 30 (6.4%) recalled being bitten by a triatomine. Moreover, 169/472 respondents (35.8%) indicated familiarity with CD and 26.2% indicated awareness of transmission routes, with 98.4% of this group signalling the vector and 1.6% indicating transfusion. Another 26 (5.5%) recognized “enlargement of the heart” as a symptom of CD. In Bolivia, 54% of participants had lived in a rural area and 37.5% had worked in a rural area. Additionally, slightly over half of respondents (50.4%) had lived in houses with mud walls; others had lived in homes of plastered (36.5%) or unplastered (6.1%) brick and cement or wood (0.9%). Furthermore, 37.1% of respondents had practiced hunting and 40% had handled meat from wild game, but these factors were not significantly associated with knowledge of the vector.

**Table 2.** Exposure to Risk Factors and Awareness of Chagas Disease and Triatomines (Vinchucas), Bolivian Migrants in São Paulo

| Factor | Total sample<br>N=472<br>N (%) | Aware of<br>Chagas disease<br>N=169<br>N (%) | p value | Aware of<br>vinchucas<br>N=225<br>N (%) | p value |
| --- | --- | --- | --- | --- | --- |
| Gender |  |  |  |  |  |
| Female | 255 (54.0) | 78 (30.6) | 0.010 | 112 (43.9) | 0.077 |
| Male | 217 (46.0) | 91 (41.9) |  | 113 (52.1) |  |
| Age |  |  |  |  |  |
| ≥30 years old | 221 (46.8) | 87 (39.4) | 0.130 | 110 (49.8) | 0.390 |
| <30 years old | 251 (53.2) | 82 (32.7) |  | 115 (45.8) |  |
| Department of birth |  |  |  |  |  |
| La Paz | 358 (75.8) | 109 (30.4) | <0.001 | 149 (41.6) | <0.001 |
| Other | 114 (24.2) | 60 (52.6) |  | 76 (66.7) |  |
| Education |  |  |  |  |  |
| < high school | 152 (32.2) | 46 (30.3) | 0.083 | 54 (35.5) | <0.001 |
| High school graduate | 320 (67.8) | 123 (38.4) |  | 171 (53.4) |  |
| Lived in a rural area of Bolívia |  |  |  |  |  |
| Yes | 277 (58.7) | 100 (36.1) | 0.945 | 145 (64.4) | 0.022 |
| No | 190 (40.3) | 68 (35.8) |  | 79 (41.6) |  |
| Worked in a rural area of Bolivia |  |  |  |  |  |
| Yes | 209 (45.2) | 74 (35.4) | 0.995 | 112 (53.6) | 0.018 |
| No | 254 (54.8) | 90 (35.4) |  | 108 (42.5) |  |
| Had a relative with Chagas disease |  |  |  |  |  |
| Yes | 48 (11.7) | 35 (72.9) | <0.001 | 35 (72.9) | 0.001 |
| No | 362 (88.3) | 123 (34.0) |  | 169 (46.7) |  |
| Received a blood transfusion |  |  |  |  |  |
| Yes | 27 (5.7) | 11 (40.7) | 0.582 | 12 (44.4) | 0.730 |
| No | 445 (94.3) | 158 (35.5) |  | 213 (47.9) |  |
| Type of housing in Bolivia |  |  |  |  |  |
| Mud or wood walls | 238 (50.4) | 71 (29.8) | 0.006 | 108 (45.4) | 0.315 |
| Brick or cement | 234 (49.6) | 98 (41.9) |  | 117 (50.0) |  |
| Saw a *vinchuca* in their home |  |  |  |  |  |
| Yes | 106 (22.5) | 71 (67.0) | <0.001 | 101 (95.3) | <0.001 |
| No | 270 | 85 (31.5) |  | 102 (37.8) |  |
| Remembers being bitten by a *vinchuca* |  |  |  |  |  |
| Yes | 30 (6.4) | 21 (70.0) | <0.001 | 28 (93.3) | <0.001 |
| No | 285 | 105 (36.8) |  | 137 (48.1) |  |
| Went hunting in Bolivia |  |  |  |  |  |
| Yes | 175 (37.1) | 56 (32.0) | 0.186 | 82 (46.9) | 0.786 |
| No | 297 (62.9) | 113 (38.0) |  | 143 (48.1) |  |

| Handled game in Bolivia |  |  |  |  |  |
| --- | --- | --- | --- | --- | --- |
| **Yes** | 189 (40.0) | 63 (33.3) | 0.360 | 92 (48.7) | 0.720 |
| **No** | 283 (60.0) | 106 (37.5) |  | 133 (47.0) |  |

Male gender, being born outside of the department of La Paz, having a relative with CD, not having lived in a house with mud walls, and having seen or been bitten by a vinchuca were all significantly associated with awareness of CD in the univariate analysis. Having lived in a rural area of Bolivia or in a department other than La Paz, having a relative with CD, having a high school education, and having seen or been bitten by a vinchuca were significantly associated with knowledge of the vector. In a multivariate logistic regression of the different risk factors plus age, education, gender, and income, only knowledge of the vector and having a relative with CD were significantly associated with awareness of CD (Table 3).

**Table 3 –.**
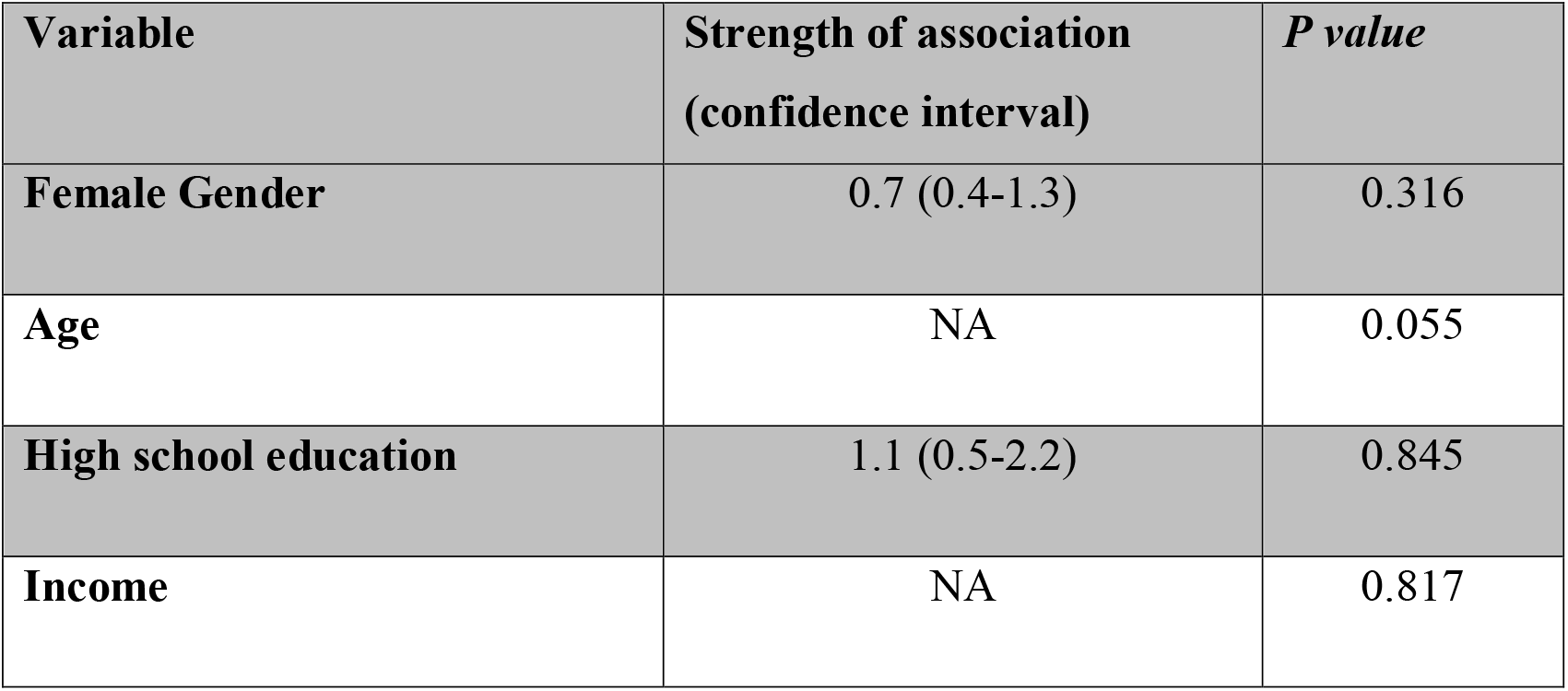
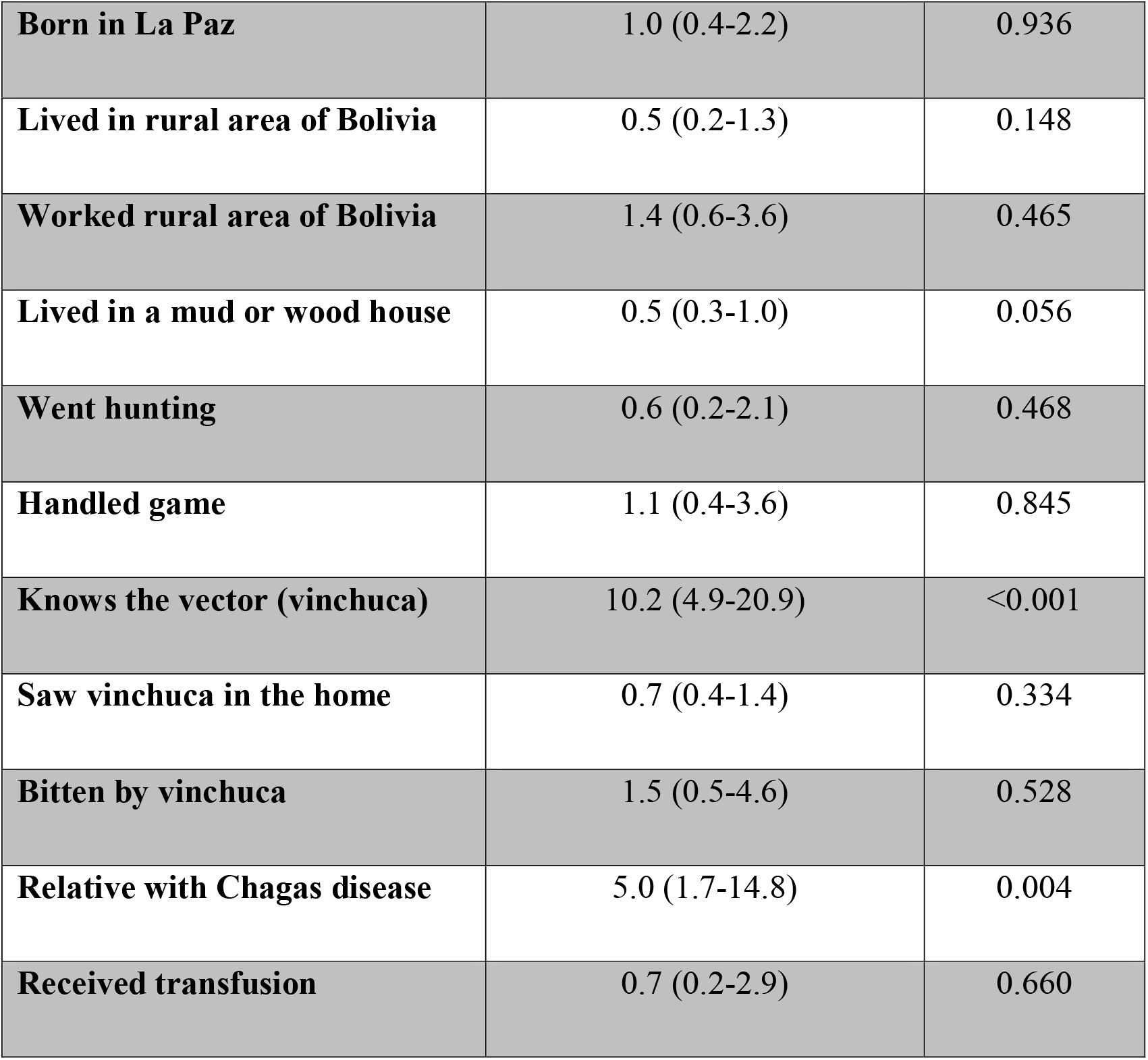
Multivariable logistic regression of factors associated with awareness of Chagas Disease

Twenty-five participants had positive serology for *T. cruzi*; the CD prevalence in the sample was 5.3%. Of these, 19 were female. Four of the seropositive individuals indicated they had previously donated blood (two in Bolivia, one in Brazil, and one did not indicate where), whereas one had received a transfusion. Of the 255 females in the sample, only 31 (12.2%) were age 40 or older, 246 (96.5%) were of childbearing age (10-49 years old) and 199 (78%) had children. While 73.7% of the mothers had had a prenatal exam (71.8% in Brazil and 26.6% in Bolivia), in only 5.3% of these cases was CD testing included, and only 1.5% indicated their infants had been tested for CD. Of the women who had had prenatal exams, >90% were aware of and had used the SUS for this service; 1.5% had coverage from private insurance and 2.6% paid out of pocket.

## Discussion

The epidemiology of CD in the twenty first century has been framed in part by the social and economic forces driving globalization [4–5]. Our study provides insight on South-South migration, which has important implications for the distribution of CD and access to healthcare. The majority of respondents were not familiar with CD. Nonetheless, the CD prevalence in the sample was 5.3%, and many had been exposed to risk factors including triatomines and/or housing susceptible to triatomine infestation, as was discussed in an earlier study [19]. Although not confirmed in multivariate analysis in our sample, perhaps due to different region of origin. there was a trend to lower awareness of CD among individuals who had lived in houses with mud or wood walls in Santa Cruz de La Sierra, Bolivia, who are expected to have a higher risk for CD [25]. One possible explanation is that socioeconomic factors have enabled greater access to health information on CD for people who live in housing with plastered walls.

Other studies have noted varying levels of CD awareness among different immigrant populations. Ramos and colleagues found that 63% of Bolivian immigrants in a sample in Elche, Spain had heard of CD [26]. In another study in Valencia, Spain of 96 Bolivian immigrants, 73 (76%) had some awareness of CD, and most recognized it had an asymptomatic phase [27] In a sample of 43 Bolivians in Munich, Germany, 30 (69.8%) indicated they had previously received information about CD, but the majority considered they had limited familiarity with transmission and symptoms, and >90% had not previously been tested [28].

Bolivians in our sample exhibited lower familiarity with CD than in these studies. This could be explained by respondents’ relative youth; many were born prior to the widespread transmission interruption campaign at the beginning of the millenia [29]. By comparison, knowledge of CD was significantly higher in older individuals among Bolivians in Valencia [27]. Another possibility is that the bulk of respondents in our study were from La Paz, which has lower levels of CD compared to other departments and was certified free of vector transmission by *Triatoma infestans* [30].

However, in a sample of Latin American immigrants in Los Angeles, most of whom were from Mexico and Central America, only 14% had previously heard of CD [11]. Similarly, among a sample of migrants at the Guatemala-Mexico border, 80% had not previously heard of CD [12].

Twenty-five participants had positive serology for *T. cruzi,* as was reported in a previous article [19], which found that seropositive respondents were significantly more likely to have knowledge of CD and the vector. Most testing positive were females, yet only 5.3% of mothers had received CD testing as part of their prenatal care. Four of the seropositive individuals indicated they had previously donated blood, whereas one had received a transfusion. Of the four donors, two indicated they had given blood in Bolivia and one in Brazil (the fourth did not specify a location).

It is essential to note that low awareness is but one of numerous barriers addressing access to diagnosis and treatment for immigrants and other groups afflicted by CD. Awareness can only be understood and addressed in terms of its interrelationship with socioeconomic inequalities, gaps in the public health response to CD (often a function of political decisions impacting public funding and concerning which populations and health issues ot prioritize), and navigation of cultural and linguistic differences. Farmer [31] points out that entrenched global political and economic structures largely shape disease epidemiology; vulnerable groups, who bear the heaviest burden, also have the least access to healthcare resources and the strongest limitations on their agency São Paulo has attracted large numbers of migrants in search of employment; this population faces significant barriers to accessing healthcare [32–35]. Bolivians in São Paulo are primarily young, and while most have attained a high school education, their income level remains low. Concentrated in São Paulo’s garment industry, they often difficult living and working conditions [34–35], which exacerbate health risks, especially for tuberculosis and other infectious diseases [32]. Although Brazilian law favors universal healthcare, undocumented immigrants, still face substantial bureaucratic difficulties in obtaining services from the SUS [36].

Moreover, awareness of CD is also a key concern within the local health system; vector transmission was interrupted in the state of São Paulo in the late 1960s [15, 37], and many providers in the SUS are unfamiliar with CD [38]. While Bolivians are a population at particular risk of CD, systematic screening, even for women in prenatal care, is not widely implemented. In our study, only 5.3% of women received CD screening as part of their prenatal care, which represents a missed opportunity to halt vertical transmission and prevent a lifelong disease. Further, the bulk of our sample was under 40, and could therefore still benefit, if seropositive, from timely etiological treatment to prevent future complications from chronic CD. Such treatment also acts as an effective means of eliminating vertical transmission [8, 39, 40].

Education campaigns to improve awareness of CD, which are culturally and linguistically tailored to the Bolivian population of Sao Paulo, should be accompanied by intensified training and capacitation of primary care personnel at facilities which see large numbers of Bolivian patients. Diagnosis with CD can be emotionally devastating and may entail stigmatization, which can actually discourage patients from seeking testing. Ideally, healthcare for CD would take a holistic approach, addressing not only the disease but its social determinants and emotional consequences [41]. Finally, care should be taken to heighten awareness in a way that does not create an unfavorable image of Bolivians and/or immigrants.

## Acknowledgments

The Drugs for Neglected Diseases *initiative* (DND*i*) is grateful to its donors, public and private, who have provided funding to DND*i* since its inception in 2003. A full list of DND*i*’s donors can be found at http://www.dndi.org/donors/donors/.

## Conflict of interest

The authors declare no conflict of interest.

^ Members of the Chagas Disease Research Group: Primary Care and Immigration Cássio Silveira, Fernando Mussa Abujamnra Aith, Lia Maria Brito da Silva, Noêmia Barbosa Carvalho, Célia Regina Furuchó, Camila G. Satolo, Pedro Albajar-Viñas, Magda Maya Atala, Vera Lúcia Teixeira de Freitas, Rosário Quiroga Ferrufino, Luzia Martinelli, Sonia Regina de Almeida.

## Supporting information

**S1 File. Datas Codification.**

**S2 File. Income.**

**S3 File. General Population.**

